# *PREDICT:* Advancing Accurate Gene Expression Prediction and Motif Identification in Plant Stress Responses

**DOI:** 10.1101/2024.03.28.587275

**Authors:** Lavakau Thalimaraw, Wei Xiong Henry Eo, Ming-Jung Liu, Ting-Ying Wu

## Abstract

Cells respond to environmental stimuli through transcriptional responses, orchestrated by transcription factors (TFs) that interpret the gene *cis*-regulatory DNA sequences, determining gene expression dynamics timing and locations. Diversification in TFs and *cis*-regulatory element (CRE) interactions result in unique gene regulatory networks (GRNs) that underpin plant adaptation. A primary challenge is identifying Transcription Factor Binding Motifs (TFBMs) for temporal and condition-specific gene expressions in plants. While the Multiple EM for Motif Elicitation (MEME) suite identifies stress-responsive CREs in Arabidopsis, its predictive power for gene expression remains uncertain. Alternatively, the *k*-mer approach identifies CRE sites and consensus TF motifs, thereby improving gene expression prediction models. In this study, we harnessed the power of a *k*-mer pipeline to address sequence-to-expression prediction problems across diverse abiotic stresses, in both bryophytic and vascular plants, including monocots and dicots. Moreover, we characterized both un-gapped and gapped CREs and, coupled with GRN analyses, pinpointed key TFs within transcriptional cascades. Lastly, we developed the <u>P</u>redictive <u>R</u>egulatory <u>E</u>lement <u>D</u>atabase for Identifying *<u>C</u>is*-regulatory elements and <u>T</u>ranscription factors (PREDICT), a web tool for efficient *k*-mer identification. This advancement will enrich our understanding of the *cis*-regulatory code landscape that shapes gene regulation in plant adaptation. PREDICT web tool is available at [http://predict.southerngenomics.org/kmers/kmers.php].

## Introduction

The fluctuating environments lead to the phenotypic diversities of crops, aiding their adaptation to various conditions and contributing to economic improvement. Previous studies have shown that transcriptional regulatory mechanisms play a crucial role in regulating gene expressions, which in turn contributes to phenotypic diversity across species (1). The transcription regulatory mechanism comprises at least two components: transcription factors (TFs) and *cis*-regulatory sites. A TF typically recognizes multiple, slightly varied *cis*-regulatory sites collectively referred to as a *cis*-regulatory element (CRE), representing the binding specificity of TFs (2). Consequently, variations in gene expression may arise from differences in the *cis*-regulatory sites on genes and/or the TF binding specificities to genes. Such variation serves as the driving force for the expansion of gene regulatory networks (GRNs) throughout the evolution of land plants for climate adaptation (3, 4). Therefore, developing a method to identify Transcription Factor Binding Motifs (TFBMs) for building the GRNs and elucidating transcriptional regulatory cascades becomes a significant question. The Multiple EM for Motif Elicitation (MEME) suite is widely used for *de novo* motif discovery, requiring no prior information on TFBMs, and has successfully identified several known stress-responsive CREs among stress-regulated genes in Arabidopsis (5, 6). This suggests the application of the MEME suite in identifying TF binding sites and subsequently summarizing them into the consensus TF motifs in plants. However, although the MEME suite is a popular approach for elucidating several TFBMs, the extent to which this pipeline can efficiently uncover the sets of TFBMs that well predict and explain plant gene expression regulation remains unclear (1).

We previously introduced a *k*-mer pipeline to efficiently and simply identify putative *cis*-regulatory elements (pCREs) from stress-response genes in tomatoes, Arabidopsis, and Marchantia (1, 3). These *k-mer-identified* pCREs were well predictive of gene expressions and some of them are similar to known TFBMs, suggesting they are authentic *cis*-regulatory elements. Here we expanded *k*-mer analyses to multiple stresses across bryophytic, monocot, and dicot plants and further assessed its application in identifying both un-gapped and gapped TFBMs. Coupled with GRN analyses, we further explore the pCREs likely bound by hub TFs to uncover transcriptional regulatory cascades. Compared to the MEME suite, the *k*-mer approach not only achieves this goal of identifying TFBMs, but also allows us to further map motif sites onto genes, predict gene expressions, and identify key TFs in GRNs. Together, our study developed a *k*-mer pipeline suited for gene expression prediction and built a web tool (PREDICT) for the easy and rapid identification of k-mer and the likely bound TFs for the general public.

## Result and Discussion

### *k*-mer pipeline effectively searches for motifs to predict gene expression in response to stress conditions

In our study of the evolution of transcriptional mechanisms and their association with differential stress tolerance across plant species, we re-analyzed publicly available RNA-seq datasets from different plant species subjected to various types of abiotic stress (1, 3, 4, 7–10) (Fig. S1A & S1B, Supplemental file 1). Subsequently, we clustered these stress-regulated genes based on their distinct expression patterns. These datasets included responses to wounding, salt, and high temperature stresses in two phylogenetically distant species, *Marchantia polymorpha,* and *Arabidopsis thaliana*, as well as a crop, *Solanum lycopersicum*. We utilized a *k*-mer (an oligomer with a length of *k* ≥ 5 base-pairs) pipeline to identify significantly enriched sequences among genes in different clusters (Fig. 1A. See Methods). We then compared the motifs identified from the *k*-mer pipeline with those identified using the MEME pipeline, a classic and well-recognized approach for discovering *cis*-elements (Fig. 1A. See Methods). We observed that the *k*-mer pipeline tends to identify a larger number of sequence motifs compared to the MEME pipeline (∼30-100 from *k*-mers vs. ∼10 from MEME, respectively, Fig. S1C, Supplemental file 1).

**Figure 1.**
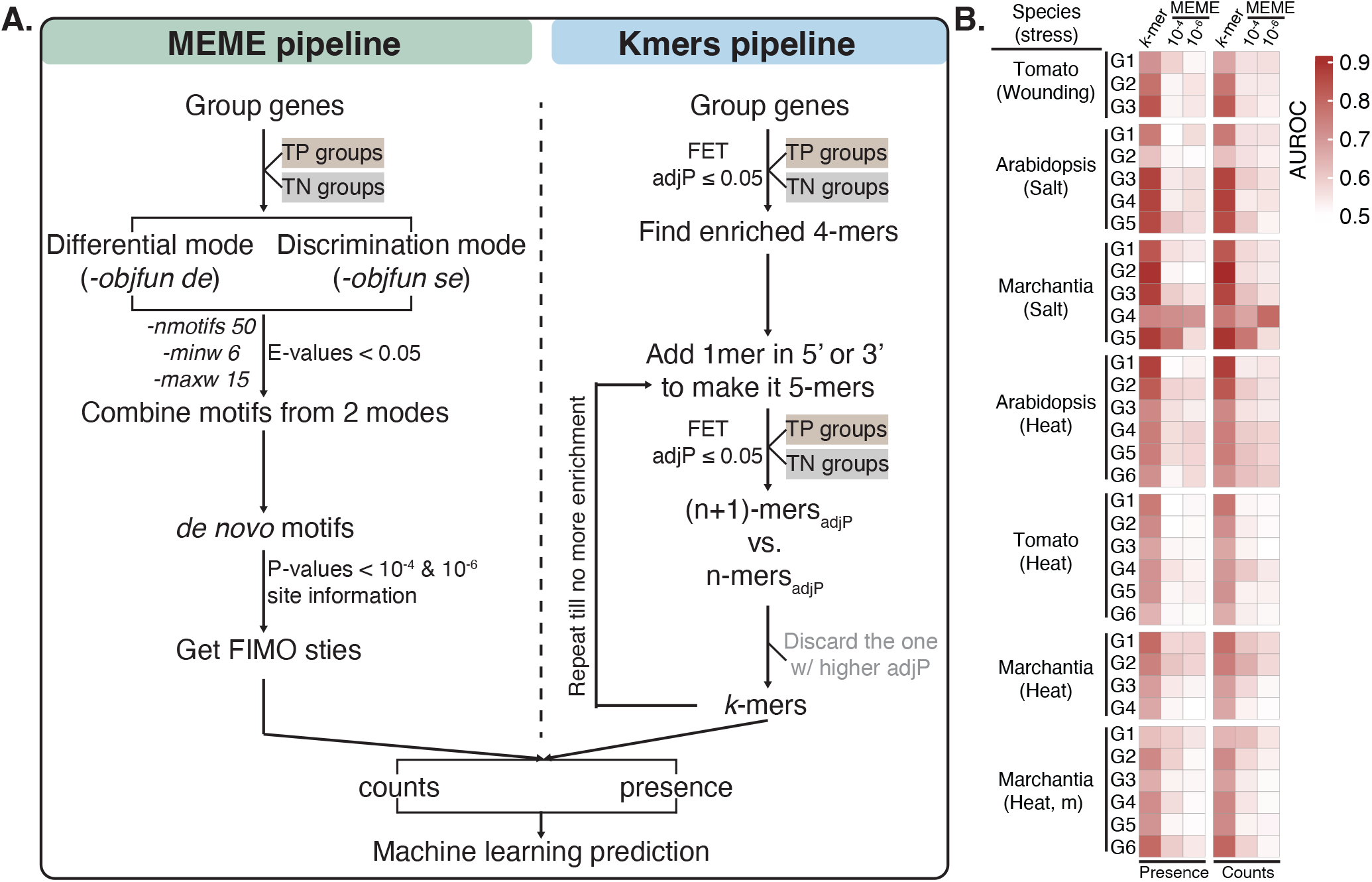
The *k*-mer pipeline outperforms the MEME pipeline for predicting stress-responsive gene expression. A. The *k*-mer and MEME pipeline workflows for motif identification and machine learning (ML) prediction. B. Heatmaps show ML performance (AUROC scores) of stress response prediction using *k*-mer or MEME pipelines for gene clusters with distinct regulatory patterns upon wounding in tomato, salt stress in Arabidopsis, and Marchantia, and heat stress in Marchantia, Arabidopsis, and tomato. The thresholds of 10^-4^ and 10^-6^ were applied to infer a motif site of MEME-derived pCREs on genes (See Material and Methods). The AUROC scores are the mean values of 10-fold cross-validation. The full prediction results are shown in Supplemental file S1.

To assess the relationship between the stress response and the short DNA sequences identified through the *k*-mer and MEME approaches, respectively, we evaluated how effectively these enriched short sequences could explain and predict the stress response in each gene cluster. We utilized machine-learning (ML) algorithms (See Methods) to predict stress-responsive genes based on the identified short sequences. In each cluster, the stress-response prediction model utilizing *k*-mer-derived short sequences outperformed the model that used MEME-derived sequences. This was indicated by AUROC scores ranging from 0.6-0.9 from the *k*-mer pipeline and 0.5-0.6 for the MEME pipeline with few exceptions with the salt stress in Marchantia G4 (Fig. 1B, Supplemental Fig. S1C, Supplemental file S2). These results showed that the short DNA sequences identified through the *k*-mer approaches were likely the putative *cis*-regulatory elements (pCREs). Furthermore, the *k*-mer pipeline is more efficient in identifying pCREs for stress-responsive genes across various stress types and plant species, suggesting the efficiency and generality of the *k*-mer pipeline in detecting cis-elements. While the MEME suite has been widely used and successful in identifying CREs for tissue- and/or stress-responsive gene expressions in Arabidopsis (6, 11), we observed its subpar performance in predicting the stress response of genes in this study. One possible reason could be the loss of sequence information of motifs generated by the MEME suite. For each motif, MEME converts all possible motif sites into position weight matrices (PWM), which can result in a loss of specific sequence information. Consequently, when mapping motifs back to the genome, this could lead to high false positive rates in identifying *cis*-element sites, resulting in decreased prediction performance.

### *k*-mer pipeline identifies the *cis*-regulatory elements near TSSs

Next, we investigated whether the pCREs identified in stress-responsive clusters are predominantly located around transcriptional start sites (TSSs), a regulatory region known for TF binding (12). We examined the site distribution of pCREs identified from the *k*-mer and MEME pipelines in the region extending from 1 kb upstream to 0.5 kb downstream of TSSs. The site distributions from the *k*-mer pipeline were primarily located near TSSs, while those from the MEME pipeline were dispersed randomly along the promoter sequence regions in all stress clusters (Fig. 2). This pattern was particularly pronounced in the case of wounding and salt stress (Fig. 2A & 2B), while the motif distributions in HS were more scattered, especially in Marchantia (Fig. 2C & 2D, Supplemental Fig. S1D). For example, the k-mer pipeline-generated pCREs similar to the motifs of HSF, NAC, and MYB were located near TSSs, consistent with evidence from ChIP-seq data in previous studies (Fig. 2E-2G, Supplemental Fig. S1E) (8, 13, 14). These findings suggested that the information extracted from the *k*-mer pipeline could provide an alternative method for searching for functional TF motifs on the core promoters.

**Figure 2.**
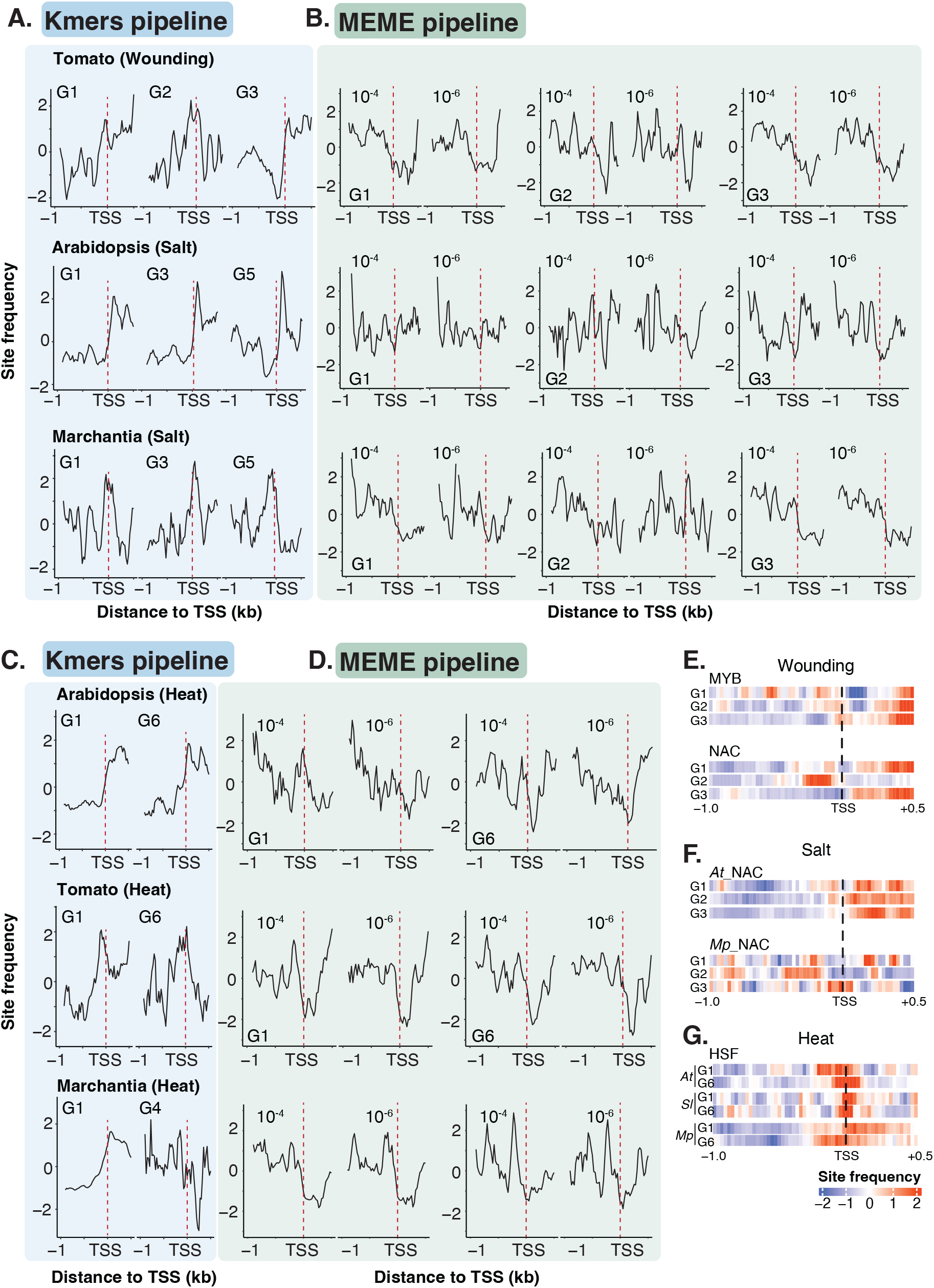
The *k*-mer pipeline identifies motifs that are preferentially located near transcriptional starting sites (TSSs). A-B. Distribution of predicted cis-regulatory elements (pCREs) identified by the *k*-mer pipeline (A) or MEME pipeline (B) during wound stress in tomato, and salt stress in Arabidopsis and Marchantia, respectively. C-D. Similar distributions as in (A) and (B), but for heat stress in Marchantia, Arabidopsis, and tomato. E-G. Heatmaps show the site distributions of k-mer pipeline-derived pCREs that are similar to known TFBMs in response to wounding (E), salt (F), and heat (G) stress, respectively. Site frequency represents the Log2 enrichment of pCRE sites between TPs and TNs and is calculated within a range from 1 kb upstream to 0.5 kb downstream of TSSs using a sliding window of 100 bp and a step size of 25 bp. The mean site frequency values for all pCREs in each cluster are displayed.

On the other hand, studies of *cis*-elements that regulate target genes in a location- and orientation-independent manner, known as enhancers, are just beginning because they often relate to transcription burst frequency (i.e., the probability of transcription activation) rather than the magnitude of mRNA synthesis (i.e., gene expression) (15, 16). Similarly, identifying regulatory sequences that silence gene expression, known as silencers, has been challenging due to the difficulty of classifying the genomic elements (17, 18). Nevertheless, the flexibility of the *k*-mer pipeline can facilitate the identification of those context-specific features within the genome.

### *k*-mer pipeline identifies the TFs and their known TFBMs in an expression-specific manner

Given that we identified only approximately 10 motifs using the MEME pipeline and with poor prediction performances (Fig. 1), we asked whether stress-responsive gene expressions could be predicted based on known motif sets from plant TFs. The ARABD database with experimental validated TFBMs contains the information of ∼18000 Arabidopsis TFBMs from 529 TFs (19) was applied here. We examined the enriched TFBMs and their corresponding binding sites via Analysis of Motif Enrichment (AME; See Method) approaches for the prediction of gene expression in each gene group. The AUROC scores from the AME pipeline decreased dramatically as the stringency of defining a corresponding binding site increased (i.e., the q-value cutoff decreased from 10^-2^∼10^-4^) (Fig. 3A). In addition, the AUROC scores for models using the “counts” of the motif sites on genes were higher compared to those using the “presence” dataset in the q-value cutoff of 10^-2^ and 10^-3^. However, those scores were consistent across datasets at the more stringent q-value cutoff (10^-4^). This indicated that using a less stringent cutoff in the AME pipeline tends to find similar motifs repeatedly in a set of promoters, leading to overfitting in gene expression prediction. In contrast, the *k*-mer pipeline identified specific short motifs and their perfect-matched sites on the promoters, reducing the likelihood of counting the same regions. Consequently, the predictive power when using either the “counts” or “presence” of the motif sites on genes remained consistent (Fig. 1B). These observations showed that the AME pipeline based on the DAP-seq database was not robust enough to identify models of predicting plant stress response genes.

**Figure 3.**
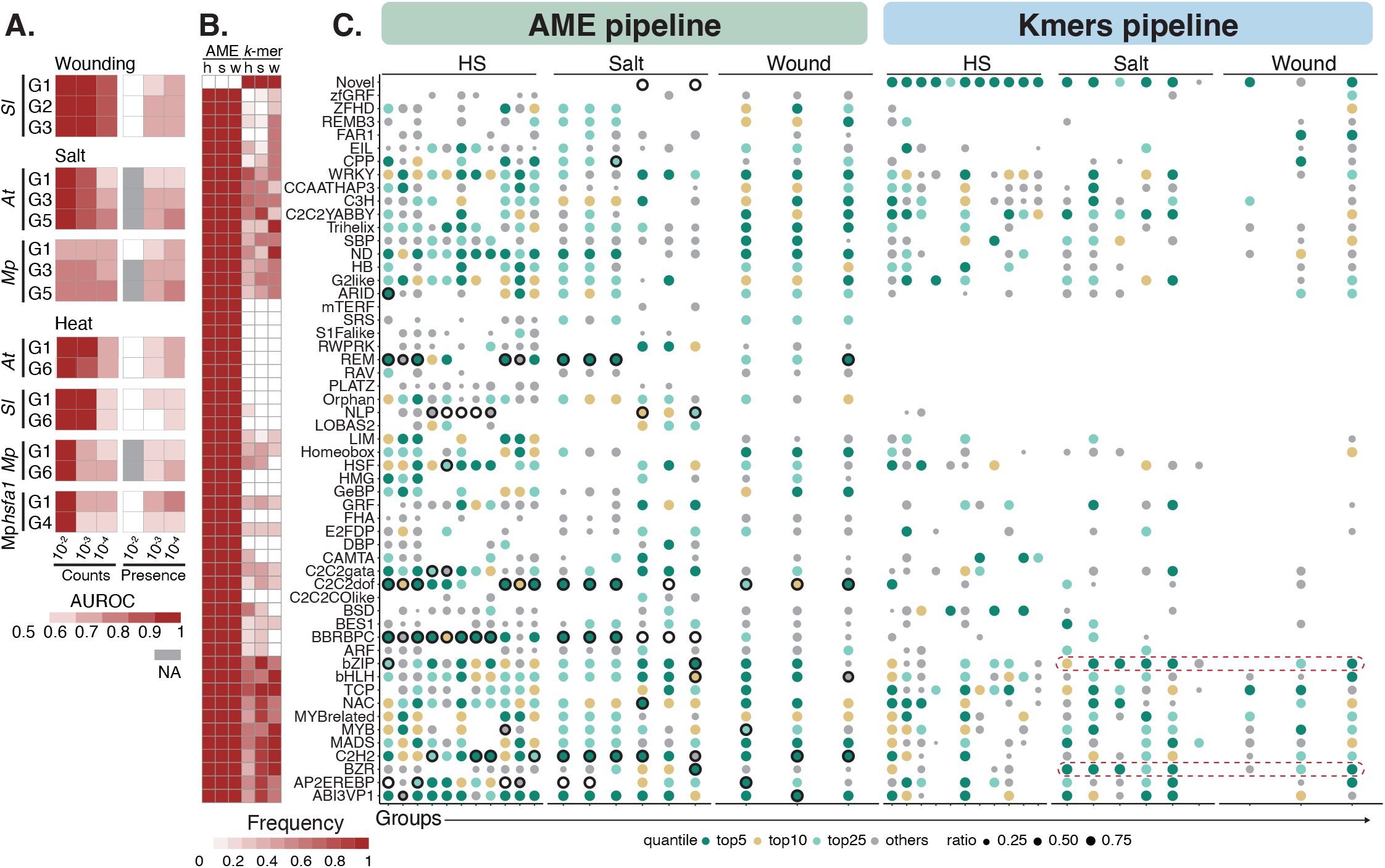
The *k*-mer pipeline identifies specific trans-regulators in various stress conditions. A. The heatmaps display prediction scores (AUROC scores) via the Analysis of Motif Enrichment (AME) pipeline (i.e, using the enriched known transcription factor binding motifs (TFBMs) filtered by different q-value cutoffs; See Material and Methods) for each stress group. B. The heatmaps summarize the frequency of transcription factor (TF) families present among clusters of each stress. Shown for the TF families inferred from known TFBMs using the AME pipeline (with q-value cutoff = 10^-2^) or *k*-mer pipeline. C. The bubble plot shows the similarity of pCREs, identified from the AME pipeline or *k*-mer pipeline, to known TFBMs in each stress group. Motif similarities to known TFBMs were calculated using the Pearson Correlation Coefficient (PCC). Quantile represents the enrichment q-values of the motifs identified from the AME or k-mer pipelines, which were standardized by quantile normalization. Motifs were grouped by TF families, and then ranked and scaled by prediction model-derived importance scores. The ratio is calculated as 1 minus the motif’s rank divided by the number of total pCREs identified in each group, ranging from 0 to 1, with higher values indicating higher importance. The highest ratio within each TF family is plotted here. The circles with black strokes indicate those TFBMs identified using the MEME pipeline. The red dashed boxes indicate the TFBMs that are specifically enriched in the salt and wounding dataset.

Furthermore, we compared the pCREs identified from the *k*-mer pipeline with TFBMs experimentally validated from the DAP-seq database (19) to determine if the identified pCREs could represent the motifs from various TF families. The pCREs were matched to known TFBMs based on their PWM similarities and were ranked by the importance score generated by the ML algorithm in the *k*-mers pipeline (Fig. 1). The TFBMs identified by the AME pipeline were present across almost all types of TF families in each cluster, suggesting those motifs may represent the general stress-responsive elements regardless of gene expression patterns (Fig. 3B&3C). In contrast, those identified by the k-mer pipeline exhibited cluster-specific patterns (Fig. 3B&3C). For example, the BZP and bZIP TF families were highly ranked and more enriched in salt and wound stress, while HSF and WRKY TF families were significantly represented in HS conditions. Additionally, the *k*-mer pipeline identified novel motifs not resembling any known TFBMs, across all stress types (Fig. 3B&3C), indicating that the functions of many *cis*-regulatory codes remain unknown and await experimental validation. Together, this suggests that the motifs identified from the *k*-mer pipeline may represent stress- and species-specific TFBMs, while the AME-derived TFBMs represent the motifs that are involved in the general stress responses.

### GRNs with *k*-mers information are more informative and accurate

A transcriptional response is triggered by TFs that directly read and bind to cis-regulatory DNA sequences of genes. We further tested whether including *k*-mer information would improve the accuracy of HS GRNs. First, we constructed a global HS GRN encompassing 1959 nodes and 6090 edges and then subset this global GRN based on the confidence levels of the edges, resulting in a sub-GRN with 183 nodes and 406 edges (Fig. 4A; Supplemental Fig. S2A). Subsequently, we focused on HSFA2-centered sub-GRN as it plays an important role in HS response in plants (20). We divided the HSFA2 GRNs into three layers based on direct-linked edge information (Fig. 4A & 4B, See Method). We hypothesized that the innermost area (i.e., the 1^st^ layer) contains a higher number of genes directly targeted and bounded by a TF (targeted gene), and hence the corresponding motifs are on the promoter of targeted genes. Supporting this assumption, we found that 92% of nodes contain HSE motifs in the 1^st^ layer of the sub-GRN while only 78% of nodes contain such motifs in the 3^rd^ layer (Fig. 4C). The HSE motifs were specifically enriched in the nodes of all the layers in the network (Fig. 4D). Likewise, the GRNs with *k*-mer information included a higher ratio of benchmarked genes, such as several heat shock proteins (HSP) including sHSPs, HSP90 and DNAJ (21) (Fig. 4B, Supplemental file S3). We also observed indirect regulation (i.e., no *k*-mer information in genes) to HSFB2a, HsfB2b, HSFA7a, HSFB1 and HSFA7b (Fig. 4B).

**Figure 4.**
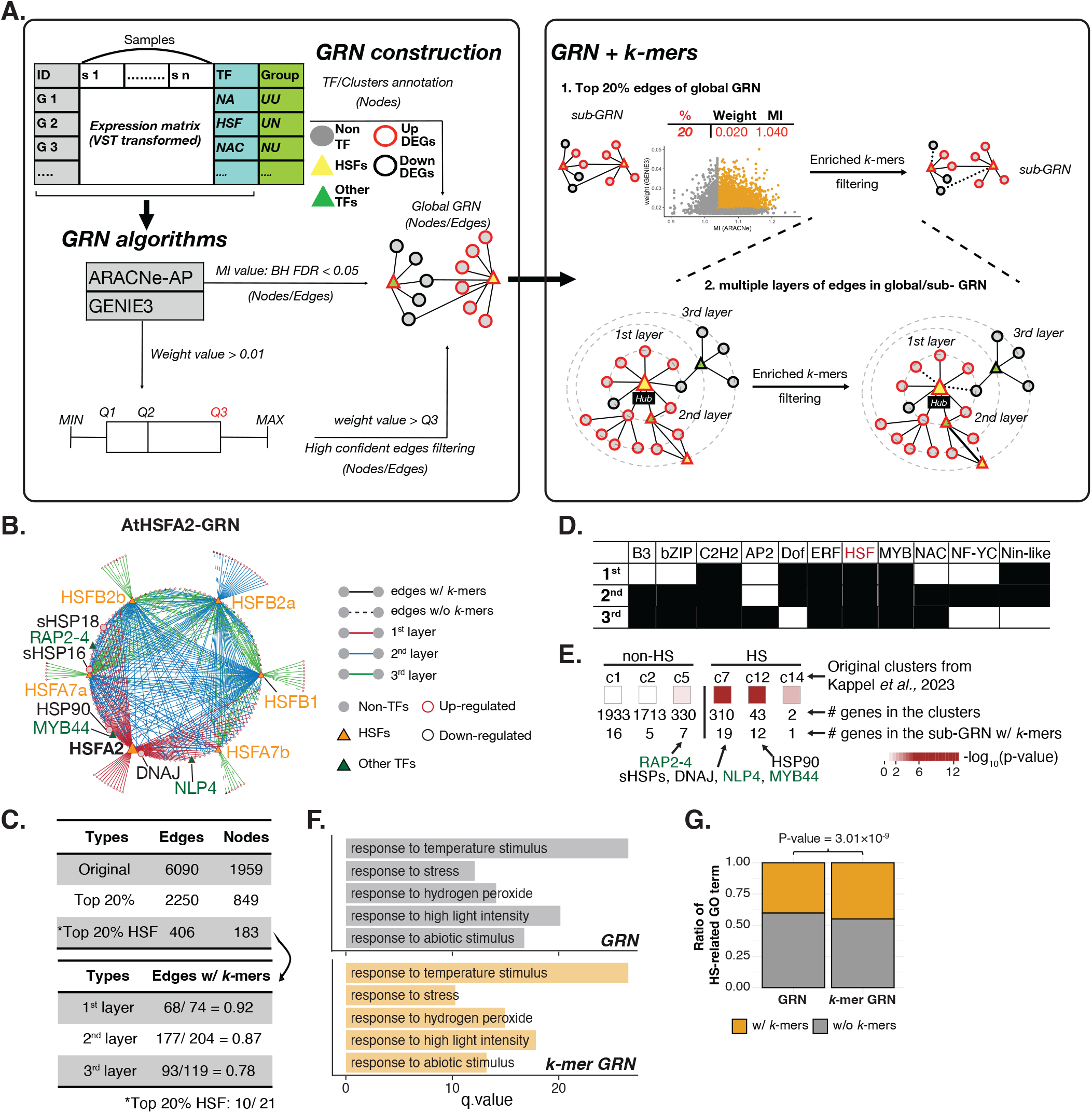
The *k*-mer pipeline improves heat-stress (HS) GRN construction. **A**. A workflow for GRN identification and inferring directly targeted genes using *k*-mer information. B. The AtHSFA2 GRN illustrates directly regulated genes (1^st^ layer) and subsequent layers, with known target genes labeled (in black). C. Summary of the number of nodes and edges with or without *k*-mer information. Shown for the HS GRN will all edges, the top 20% edges and the top 20% edges containing HSF nodes. D. The enrichment of known TFBMs in genes located in different layers of GRNs compared to the global GRN. The significance is determined by Fisher’s exact test with black cells representing *p* value <0.05. E. Number of AtHSF2 direct-binding genes in the HS-responsive and non-HS-responsive gene clusters from HSFA2 ChIP-seq dataset (22) as well as in the GRNs indicated in (B). The overrepresentation was determined by hypergeometric distribution. F. Enriched GO Biological process terms in the GRNs with or without *k*-mer information are shown, with q-value (-log_2_(adjusted p-value)) indicating the significance of the enrichments. G. Ratio of heat-related GO terms in GRNs with or without *k*-mer information, where the adjusted p-value is determined from Fisher’s exact test.

In addition to known HSP genes, the analyses of hierarchical regulations in the 1^st^ layer sub-GRN found the direct regulations from HSFA2 to NLP4, MYB44, and RAP2.4 (Fig. 4B). To determine whether the nodes with *k*-mers in the 1^st^ layer of HSFA2 sub-network were indeed the directly bound by HSFA2, we mapped them back to the public available HSFA2 ChIP-seq dataset (22). We found a higher proportion of genes significantly enriched in the clusters (c7, c12, and c14; these were directly taken from the previous publication) directly bound by HSFA2 during HS condition compared to genes in non-HS clusters (c1, c2, and c5). These genes included sHSPs, DNAJ, NLP4, and MYB44 in c7, as well as HSP90 in c12 (Fig. 4E). This observation supports the hypothesis of a direct transcriptional regulation between HSF-related TFs and other unexplored TFs. Lastly, a higher proportion of TF-targeted genes were overrepresented with HS-related GO terms in the sub-GRN that included *k*-mer information (Fig. 4F & 4G). A similar result was observed when we constructed the SlHSF2-subGRN under heat stress (Supplemental Fig. S2B-S2D).

Furthermore, when we focused on the sub-network centered by a hub gene, NAC062, which was not previously known to regulate the HS response, we observed that the enriched *k*-mers on the directly targeted genes of NAC062 were associated with NAC-binding motifs at all layers (Supplemental Fig. S2E& S2F). The GO terms derived from the *k*-mer network that resembled the NAC binding motif were significantly associated with the general abiotic stress response and response to chitin, compared to the global network (Supplemental Fig. S2G & S2H). These GO terms associated with *k*-mers that resembled the NAC binding motif differed from those associated with *k*-mers similar to HSF binding motifs (Supplemental Fig. S2I). The NAC and HSF binding motifs were specifically enriched in their 1^st^-layer GRNs (Fig. 4D and Supplemental Fig. S2F). These findings suggested that incorporating specific *k*-mers information into sub-network construction could improve the accuracy of identifying specific TF-targeted gene pairs and inferring their biological functions.

Together, this refined GRN construction workflow provided a strategic approach for the *in silico* prioritization and screening of TFs that are involved in HS response for further experimental validation. Furthermore, it is known that the integration of various data types, such as ATAC-seq or ChIP-seq, can enhance the stability of the GRN architectures (23). Applying the *k*-mer pipeline to identify motifs in these datasets could further refine the identification of direct targets, thereby untangling the complexity of transcriptional regulatory cascades under different stress conditions.

### Gapped *k*-mer pipeline identifies novel HS-responsive motifs

We further noticed that the prediction scores for HS-responsive gene clusters were lower than those for clusters associated with other types of stress, regardless of the plant species (Figs. 1B, 2C, 3B). Given that well-known HS-related motifs (Heat Shock Element, HSE) are gapped, with two wobble nucleotides between GAA and TTC (GAAnnTTC and/ or TTCnnGAA) (24), we modified our *k*-mer pipeline to enable the search to target gapped *k*-mers that are similar to HSEs (Fig. 5A, see Method). The F1 and AUROC scores for HS up-regulated clusters or the clusters with more complicated expression patterns (Fig. S1C) improved when we considered gapped *k*-mer, with up to a 10% increase in each tested species, especially in Marchantia and tomato (Fig. 5B), aligning with our hypothesis and previous observations (8, 14). Less than 10% of gapped *k*-mers matched the known TFBMs in the DAP-seq database (bottom in Fig. 5C). The gapped *k*-mers exactly matched HSEs ranked in the top 5 based on their importance scores in each cluster regardless of species (orange dots in Fig. 5C; Supplemental file S4). The gapped *k*-mers that perfectly matched or had high similarity to the HSE ranked highly in the predictions and were identified more frequently over the 10 machine learning rounds (Fig. 5D & 5E). These observations indicate their substantial contribution to the model’s accuracy. Together with the findings of the un-gapped pCREs (Figs. 1-3), these results indicate the generality and flexibility of the *k*-mer pipeline to characterize the un-gapped and gapped pCREs for diverse types of stress-related TFs.

**Figure 5.**
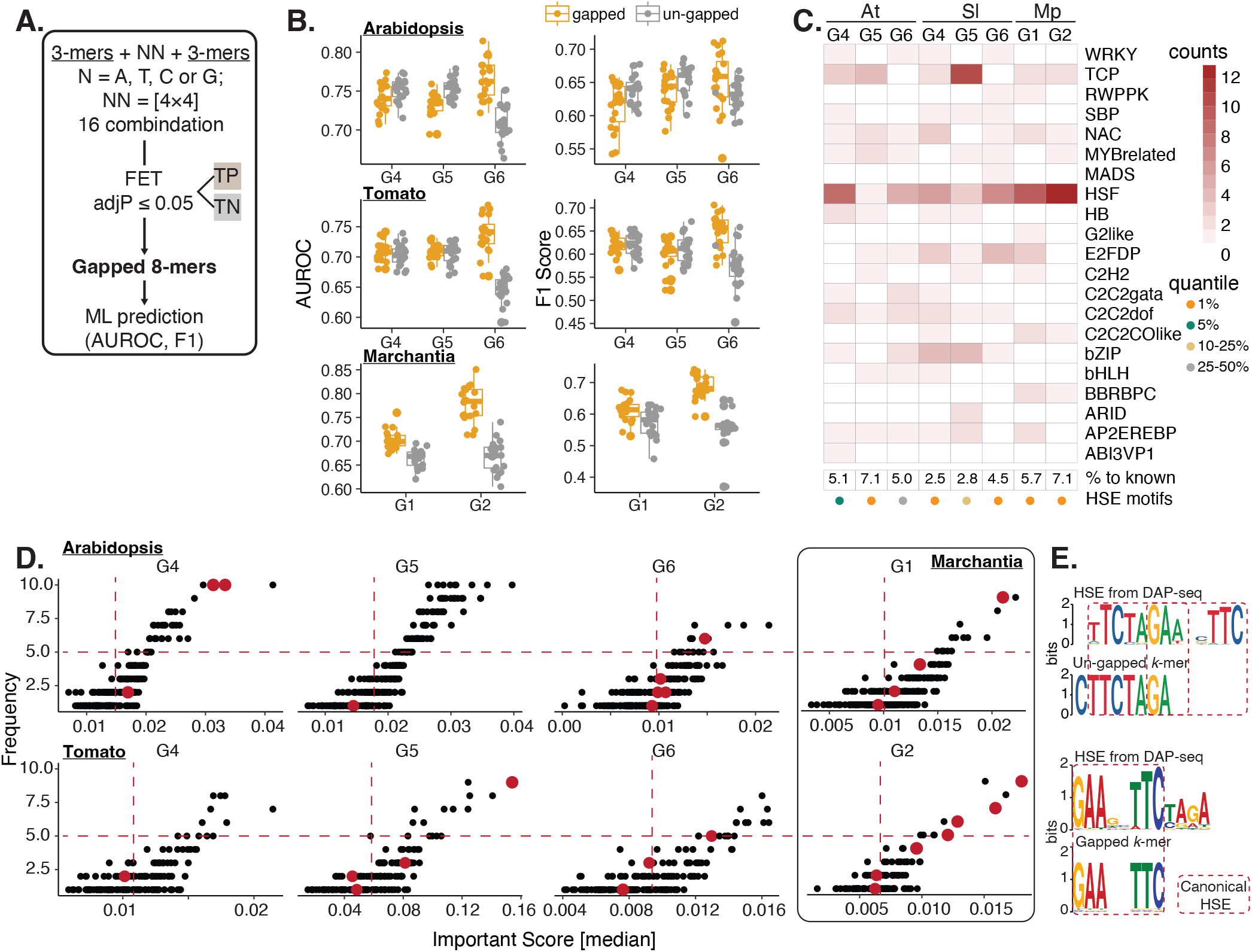
The gapped *k*-mer pipeline improves HS-response prediction in plants. A. A workflow designed for identifying gapped *k*-mers under HS conditions. NN represents the gapped sites when searching sites of a tested kmer on genes (See Material and Methods). B. The boxplots illustrate ML performance for the HS dataset in Marchantia, Arabidopsis, and tomato with gapped or non-gapped *k*-mers identification pipelines. C. A heatmap shows the counts of gapped motifs that are similar to known TFBMs. The similarity between TFBM motifs and gapped motifs and their relative importance scores, represented by quantiles, was indicated at the bottom. D. Dotplots show the frequency in 10 downsizing datasets and the importance score for each enriched gapped *k*-mer, with red dots indicating known HSE motifs. The dashed red lines represent the median value of important scores and frequency in each group. E. Representative sequence logo from un-gapped and gapped *k*-mers that matched to HSE from DAP-seq database.

### PREDICT: an easy-to-use web tool for *k*-mer identification and TFBM association

To streamline the identification and visualization of *k*-mers intuitively using our pipeline, we built a new web-based tool, <u>P</u>redictive <u>R</u>egulatory <u>E</u>lement <u>D</u>atabase for Identifying <u>C</u>is-regulatory elements and Transcription factors web tool (PREDICT), for conducting the motif search using *k*-mer pipeline, visualizing *k*-mer sites along genes and associating *k*-mers to the known TFBMs. The workflow of the PREDICT web tool can be broken down into three major steps, as shown in Fig. 6A:

**Figure 6.**
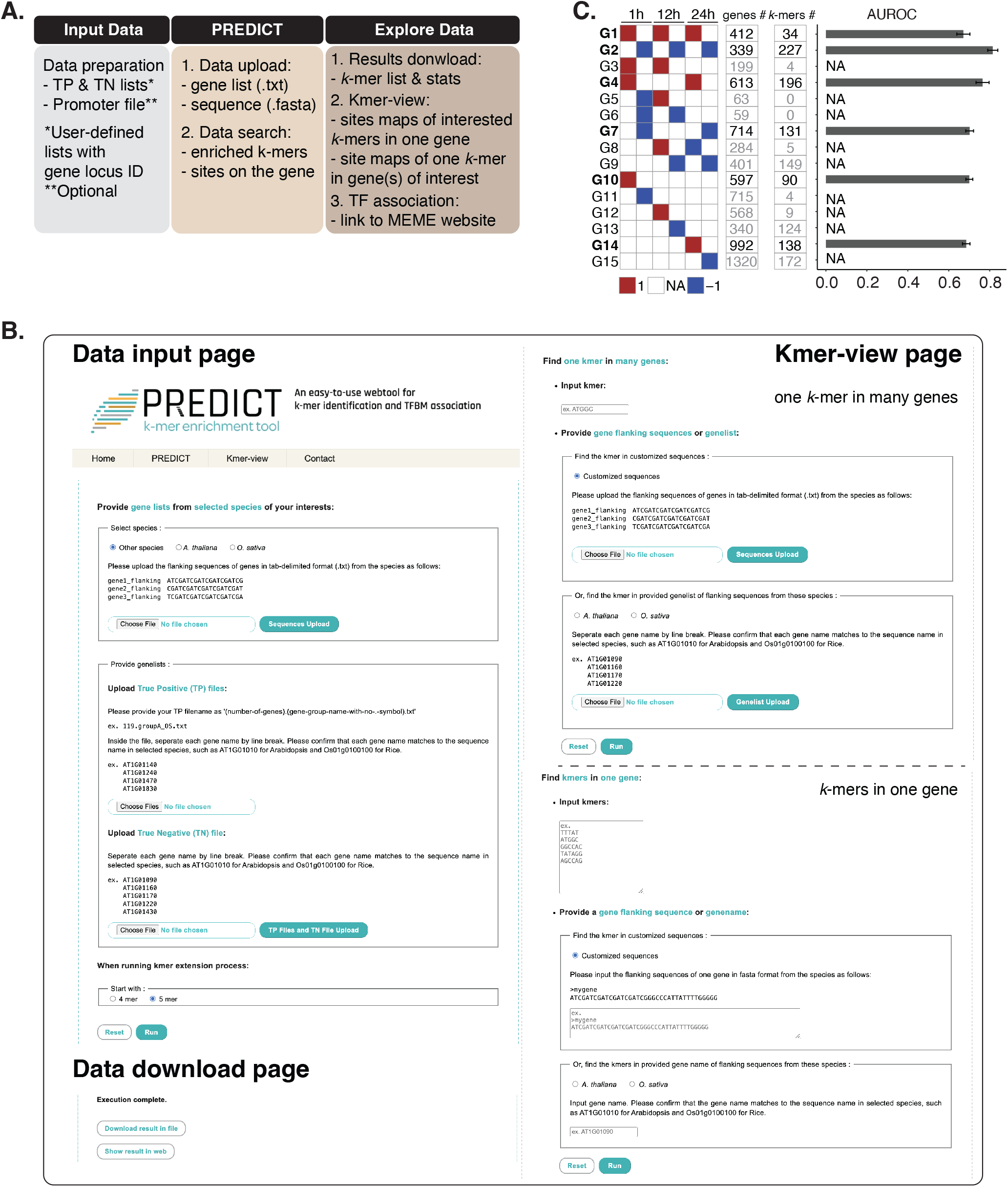
Predictive Regulatory Element Database for Identifying *Cis*-regulatory elements and Transcription factors (PREDICT) identifies enriched *k*-mers in the rice dataset under heat stress. A. Schematic representation of the major steps in the PREDICT web tool. B. Representative screenshots for PREDICT. Note that in the web tool version, the program will search starting from 4- or 5-mers until 12-mers. The PREDICT tab is for data upload and executing the program and the Kmer-view tab is for exploring the sites of one *k*-mer in multiple genes or multiple *k*-mers in one gene. C. The heat-responsive expression patterns, the gene numbers, the counts of *k*-mers, and the ML prediction performance (AUROC scores) for gene clusters identified from rice. Note that only selected groups were subjected to ML prediction due to the amount of enriched *k*-mers.

#### Data preparation

Users prepare the predefined lists of True Positive (i.e., genes with interested expression patterns) and True Negative (i.e., genes without regulatory patterns) based on the expression patterns observed in the dataset. These lists are saved into a file with a .txt extension before uploading. The users have the option to upload the promoter sequence files in a file with the tab-delimited format and with a .txt extension for the genomic regions of genes from the species of their interest, or they may choose to work with the built-in promoter files of all genes (*Arabidopsis thaliana* and *Oryza sativa cv. japonica*). We aim to expand the number of built-in genomes in the future.

#### PREDICT identification

Once users upload the prepared file to the web tool and click the “Run” button, the program initiates the identification of enriched *k*-mers. The processing time for PREDICT may vary based on the number of genes in the list, but it typically does not exceed around 2 minutes for a list of around ∼1000 genes. The results are displayed almost instantaneously after the calculations are complete.

#### Data exploration

A notification will inform users when the identification job has finished. Users can then download the lists of *k*-mers and their corresponding p-values by clicking the “Download” button with the results provided in txt file format. Users have the option to display the sites of individual *k*-mer along all genes or to view the sites of all enriched *k*-mers in a single gene. In addition, users can visit the MEME website to compare the *k*-mers with known TFBM and then potentially associate them with specific TFs.

PREDICT has built-in example files that users can load to test the programs and parameters. These examples are derived from our previous publications and analyses (Fig. 1). In addition, to demonstrate the efficacy of our web tool, we utilized an independent rice dataset at HS and drought conditions (SRP190858) for *k*-mer identification and the prediction of gene expression (Fig. 6B & C). Several *k*-mers were identified in each HS group, with counts ranging from 34-227 and with the AUROC scores varying from 0.69 to 0.82. Similarly, the *k*-mer pipeline successfully predicted the gene expressions for drought response in rice with the AUROC scores ranging from 0.77 to 0.88 (Supplemental Fig. S3). PREDICT provided an easy-to-use web service for identifying and comparing enriched *k*-mers in various stresses, which can now facilitate studies into the diversification or conservation of the *cis*-regulatory codes in stress responses in the same or different species, thereby guiding the future experiment directions. Beyond the transcriptional regulation, PREDICT also allowed us to identify the *k*-mer-derived mRNA features around the translational initiation sites (TISs), hence deepening our understanding of the mechanism of translational regulation (25). These results suggested not only that the *k*-mer pipeline can be widely applied to various research questions at different regulatory levels in stress biology, but also demonstrated the extensive utility of a PREDICT-style analysis.

## Conclusion

Over the past few years, the significance of *cis*-regulatory elements in studying the regulatory landscapes of various plant species in ever-changing environments, as well as in the field of plant synthetic biology, has been increasingly recognized and emphasized. The web tools based on known TFBMs are used to find motif sites on stress-response genes. These tools and databases have been developed to identify *cis*-regulatory elements by identifying consensus sequences based on sequence similarity of the known TFMBs (26). However, a user-friendly pipeline integrating the discovery of key *cis*-regulatory elements on pattern-specific gene clusters to the identification of precise TFBM sites along individual genes has not been established for the plant research community. In this study, we developed a PREDICT web tool with the rapid *k*-mer identification pipeline for profiling precise cis-regulatory elements on genes among different stresses and across plant species. We first demonstrate that using precise *k*-mers as progenitors can significantly enhance the prediction of gene expression under various stress conditions across different plant species. Secondly, we provide evidence that *k*-mers are valuable features for deducing stress-specific TFs. Thirdly, we illustrate that employing *k*-mers as features to refine the architecture of GRNs can lead to the generation of new hypotheses for subsequent narrowing down the candidates for experimental validation. Lastly, we have developed a web tool called PREDICT, which allows for *k*-mer searches through a graphical user interface (GUI). We believe that this all-in-one web tool will save researchers time and effort, eliminating the need to navigate various command lines for such analyses, and will enhance the understanding of the mechanisms underlying cis-regulatory elements, with potential applications in synthetic biology.

## Material and Methods

### k-mer identification pipeline

To identify putative *cis*-regulatory elements (pCREs) associated with a given stress response, a *k-*mer (oligomer with the length of k) pipeline was adopted from a previous study (1). Briefly, this pipeline identifies short sequences significantly enriched in the regulatory region of stress-response gene clusters compared to the nonresponsive gene cluster and determines adjusted p-values using Fisher’s exact test and multiple testing correction (Benjamini-Hochberg method) (27). The regulatory region is defined as the region ranging from upstream 1 kb to downstream 0.5 kb of the transcription start site (TSS) of a gene. This *k-*mer pipeline includes several steps to discover pCREs, as shown in Figure 1A. Below, we describe each step in detail.

**Step 1:** Evaluate all possible 5-mer oligomers for enrichment among genes in stress-response clusters (positive groups) versus nonresponsive clusters (negative groups). Retain only 5-mers with significant enrichment (adjusted *p-values*<0.05).

**Step 2:** Extend the sequence of the significantly enriched 5-mers from Step 1 by 1 nt in either direction, assess the enrichment, and retain those that remain significantly enriched (adjusted *p-values*<0.05). Repeat this process until no further enriched extended *k-*mers are identified. If two *k-*mers are both significantly enriched and one is a match within the other, only the one with a lower adjusted *p-value* was retained.

**Step 3:** Repeat the process from Step 1, but begin with all possible 6-mers. Combine significantly enriched 6-mers with the set of the *k-*mers identified from Step 2.

**Step 4:** Similar to step 2, but begin with a set of *k-*mers from Step 3. Finally, identify the set of *k-*mers significantly enriched in the stress response cluster and consider them as pCREs.

### Gapped k-mers identification pipeline

The gapped 8-mers sequences are designed by combining two 3-kmer oligomers separated by a 2-bp gap. A set of the gapped 8-mers is generated by randomly selecting a pair of 3-mers oligomers from all possible 3mers and inserting 2 ‘N’s in-between. The enrichment analyses were performed as described in the previous section.

### MEME pipeline for pCRE identification

Another toolkit within the motif-finding pipeline, which includes the MEME and Find Individual Motif Occurrences (FIMO) programs, was adopted to discover pCREs and identify their potential sites on genes (28). To comprehensively identify the MEME-derived pCREs, two search modes were used: Discriminative (with parameters of -mod zoops -nmotifs 50 -nostatus -minw 6 -maxw 15 -objfun classic -revcomp -markov_order 0) and Differential Enrichment (with parameters of -mod zoops -nmotifs 50 -nostatus - minw 6 -maxw 15 -objfun de -revcomp). MEME-derived pCREs with an e-value < 5e-02 identified from either search mode were included for motif site identification via FIMO.

The FIMO-based search for motif sites was conducted with parameters --bgfile --nrdb --thresh 1e-2 --parse-genomic-coord. Motif sites meeting different q-value thresholds as indicated in Figures 1 and 3 were selected for further machine learning (ML) analyses of gene expression.

### Analysis of Motif Enrichment (AME) pipeline for enriched known TFMBs searching

First, we used the FIMO program to search for the binding sites of 591 Arabidopsis TFs identified previously (19) on the genes in each group. We then determined the site enrichments for each TFBM based on using the criteria of “counts” or “presence” and “FIMO-derived q-value” ranging from 10^-2^ to 10^-4^ by one-tail Fisher’s exact test (FET). Finally, we selected the TFBMs that showed enrichment with a q-value ranging from 10^-2^ to 10^-4^ in each group, respectively, for further prediction.

### RNA-seq dataset process, true negative and positive dataset selection

Raw read counts were taken from previous studies (See Supplemental file S1). Differentially expressed genes (DEGs) were identified with the R package DESeq2 followed by comparisons to the expression level at 0 h with the threshold of |Log_2_-fold change| ≥ 1 and False Discovery Rate (FDR) ≤ 0.05 using the “lfcShrink ashr” method. Genes with |Log_2_-fold change| ≤ 0.8 and FDR > 0.05 across each group were defined as true negative (TN) datasets. The DEGs were then clusters based on their expression pattern in the case of the heat stress dataset. The other groups were directly taken from published studies (1, 3).

### Machine learning (ML) pipeline

An ML pipeline described previously was retrieved from GitHub (https://github.com/ShiuLab/ML-Pipeline; https://github.com/azodichr/ML-Pipeline/tree/master/Workshop) (29). Briefly, we used scikit-learn (v0.24.2) in Python (v3.7.0) to train and test the models. For each stress-responsive group, we split balanced data into training (70%) and testing (30%) sets and tested four classification methods: Random Forest (RF), Support Vector Machine (SVM), Logistic Regression (LR), and Gradient Boost (GB). We used ten-fold internal cross-validation to select the optimized hyperparameters. AUC-ROC score was used to select the best model for each stress-responsive group. Random downsizing addressed the imbalance between positive and negative datasets. Enriched *k*-mers maps from 10 downsizing iterations served as input for training datasets.

### Comparison of MEME and k-mer motif distribution

The comparison of motif distribution involved two steps: site retrieval using FIMO and site counting using a sliding window. Initially, sites with enriched motifs, predicted by either *k*-mer or MEME pipelines, were identified in the promoters of both positive and negative genes. In the MEME pipeline, enriched motifs were systematically scanned with FIMO using match p-value thresholds (10^-4^ or 10^-6^), and scanning templates for the specified strand. For the *k*-mer pipeline, all enriched motifs were subjected to FIMO scanning. During the subsequent site retrieval step, a sliding window approach was used to scan for motif match sites within a given DNA strain. The window size was set to 100 bp, with a 25 bp sliding interval. The average count of matched sites for each enriched motif provided values within each window. To normalize these data, z-scores were computed for the ratio of window values from positive genes to those from negative genes within the same range.

### Construction of gene regulatory network (GRN)

Normalized gene expression counts were used to construct GRNs. Time-series heat stress datasets for *Arabidopsis thaliana* and *Solanum lycopersicum* were obtained from publicly available datasets on NCBI (7–10). DEGs were identified as aforementioned. Batch effect correction was applied using ComBat-seq (30). DEGs were then categorized into different groups based on their expression at each time point in Arabidopsis and tomato datasets. *k*-mers associated with each group were identified using the *k*-mer pipeline, followed by calculating their Pearson correlation coefficients (PCC) with motifs in the Arabidopsis DAP-seq database (31). The motif family with the highest PCC was selected to represent each *k*-mer.

ARACNe-AP (32) and GENIE3 (33) were used to generate GRNs. TFs were assigned before network construction. Only edges with Bonferroni-corrected p-values below 0.05, based on mutual information (MI) values in the ARACNe-AP network, were retained. For GENIE3, a baseline edge weight of 0.01 was set, and the interquartile range (IQR) of all edge weights was determined. The third quartile (Q3) edge weight was used to filter for high-confidence subnetworks that represent regulatory interactions within GENIE3. An intersection GRN was generated by combining the outputs of both algorithms. The top 20% of edges ranked by weight and MI value were selected to form a representative subnetwork for each species. For validation, key TFs were identified as pseudo-centers to examine the top 20% subnetwork. Nodes were grouped before each GRN construction, incorporating enriched *k*-mers and motif families. Edges were labeled as “w/ *k*-mer” if the hub TFs-associated *k*-mers were present in the promoters of target nodes towards hub TFs, referred to as a “direct-linkage” relationship. Otherwise, they were labeled as “w/o *k*-mer”. The first-layer network was defined by the immediate neighboring nodes and edges connected to the selected TF whereas the second- and third-layer networks were determined by the immediate neighboring nodes and edges originating from the nodes in the first and second layers, respectively. Gene ontology analysis was performed and compared across GRNs w/ *k*-mer or w/o *k*-mer, as well as the original GRN. The GRNs were visualized by Cytoscape software v3.10.0 (34).

### Construction of PREDICT web tool

The original scripts were modified to meet the requirements for web-based automated processing. This process involved the translation of scripts from Python 2 to Python 3, as well as the adjustment of parameters for the execution loops in the pipeline. For the web application architecture, we employed the robust LAMP stack, which integrates Linux, Apache, MySQL, and PHP to ensure scalable and efficient performance. To enhance user accessibility, the final outputs are available for a text-based download option. Additionally, the results are integrated for interactive viewing on dedicated web pages.

## Data availability

PREDICT is provided as a web tool, which is freely available at [http://predict.southerngenomics.org/kmers/kmers.php]. The accession numbers of all data used in this work are described in Supplemental Fig. 1A. The data and code used to reproduce the analyses are available at https://github.com/LavakauT/ML-pipeline and https://github.com/LavakauT/KmersDiscovery_MEME.

## Acknowledgments

We thank IPMB and the AS-BCST Bioinformatics Core for high-performance computing services, Ms. Chun-Yi Chen for web tool development, Dr. Yao-Cheng Lin and Mr. Te-Chang Hsu (AS-BSCT group) for technical support, and Dr. Chia-Yi Cheng (National Taiwan University) for critical discussion for this project. The research was supported by grant NSCT 112-2311-B-001-009-MY3 to Ting-Ying Wu and NSTC 112-2311-B-001-005 and AS-CDA-111-L06 to Ming-Jung Liu.

## Author contributions

TYW and MJL conceived and designed the project, and TYW, MJL, LT, and WHE analyzed the data and contributed to the design of the web tool. LT, TYW, and MJL wrote the paper. TYW and MJL edited the paper.

## Supplemental files

**Supplemental file S1 Motifs and their enrichment derived from k-mer, AME, and MEME pipelines for each group**

**Supplemental file S2 ML performance across different stress- and species-datasets**

**Supplemental file S3 List of genes used for GRN construction**

**Supplemental file S4 List of gapped motifs identified in each dataset**

**Supplemental file S5 Motif sitemaps for pCREs from k-mer and MEME pipelines**

**Supplemental file S6 List of genes and enriched k-mers in rice heat stress dataset**

**Supplemental file S7 List of genes enriched k-mers in rice drought stress dataset**

