## Supporting_figures for "*PREDICT:* Advancing Accurate Gene Expression Prediction and Motif Identification in Plant Stress Responses"

### Supplemental Figure 1

**A.**

| | wounding | salt | heat [condition] |
|---|---|---|---|
| **Marchantia** | - | (2) | (3), [1hr, 5hr] |
| **Arabidopsis** | - | (2) | (8), (9), (10), GSE236157 [0.5hr, 1hr] |
| **Tomato** | (7) | - | (6), GSE151277 [1hr, 6hr] |

**B.**

| NN | UU | UN | NU | DD | DN | ND | UD | DU |
|---|---|---|---|---|---|---|---|---|
| 1395 | 895 | 318 | 822 | 727 | 231 | 869 | 11 | 5 |
| 2642 | 513 | 2242 | 1546 | 563 | 940 | 1882 | 213 | 88 |
| 1906 | 595 | 1371 | 976 | 466 | 1204 | 740 | 92 | 20 |

$log_2FC \geq 1$ NA $log_2FC \leq 1$

**C.**

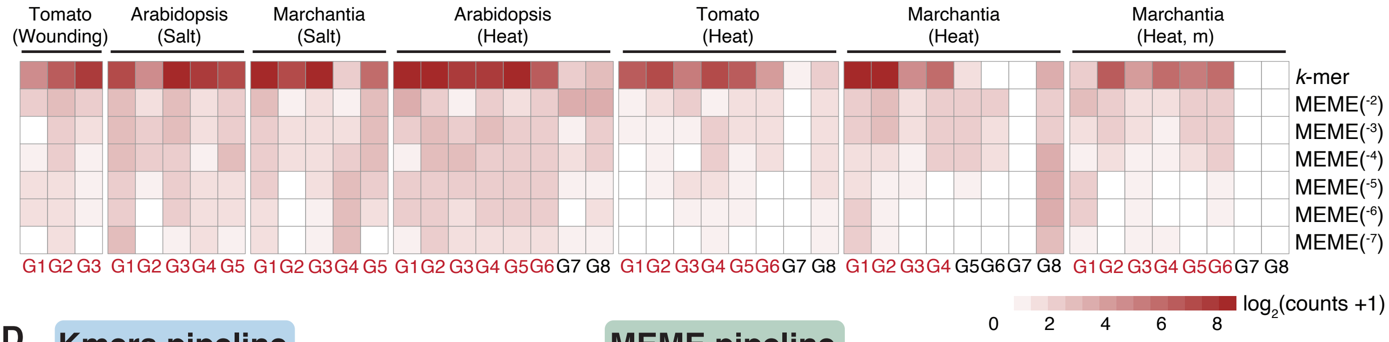


**D.**

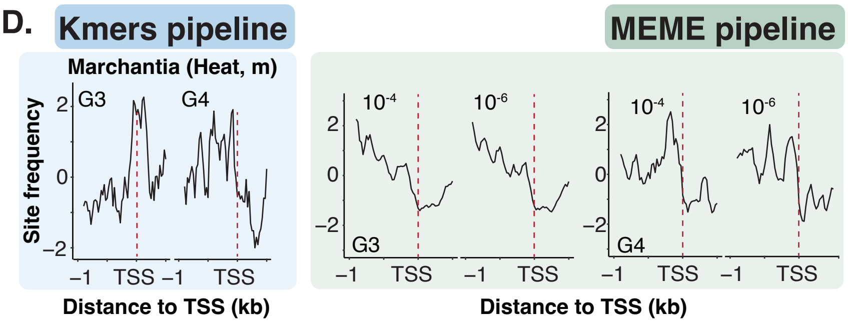


**E.**

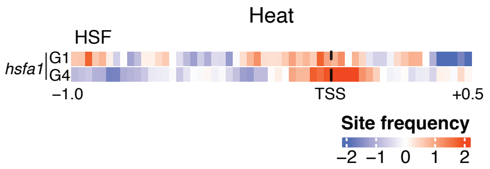

### Supplemental Figure 2

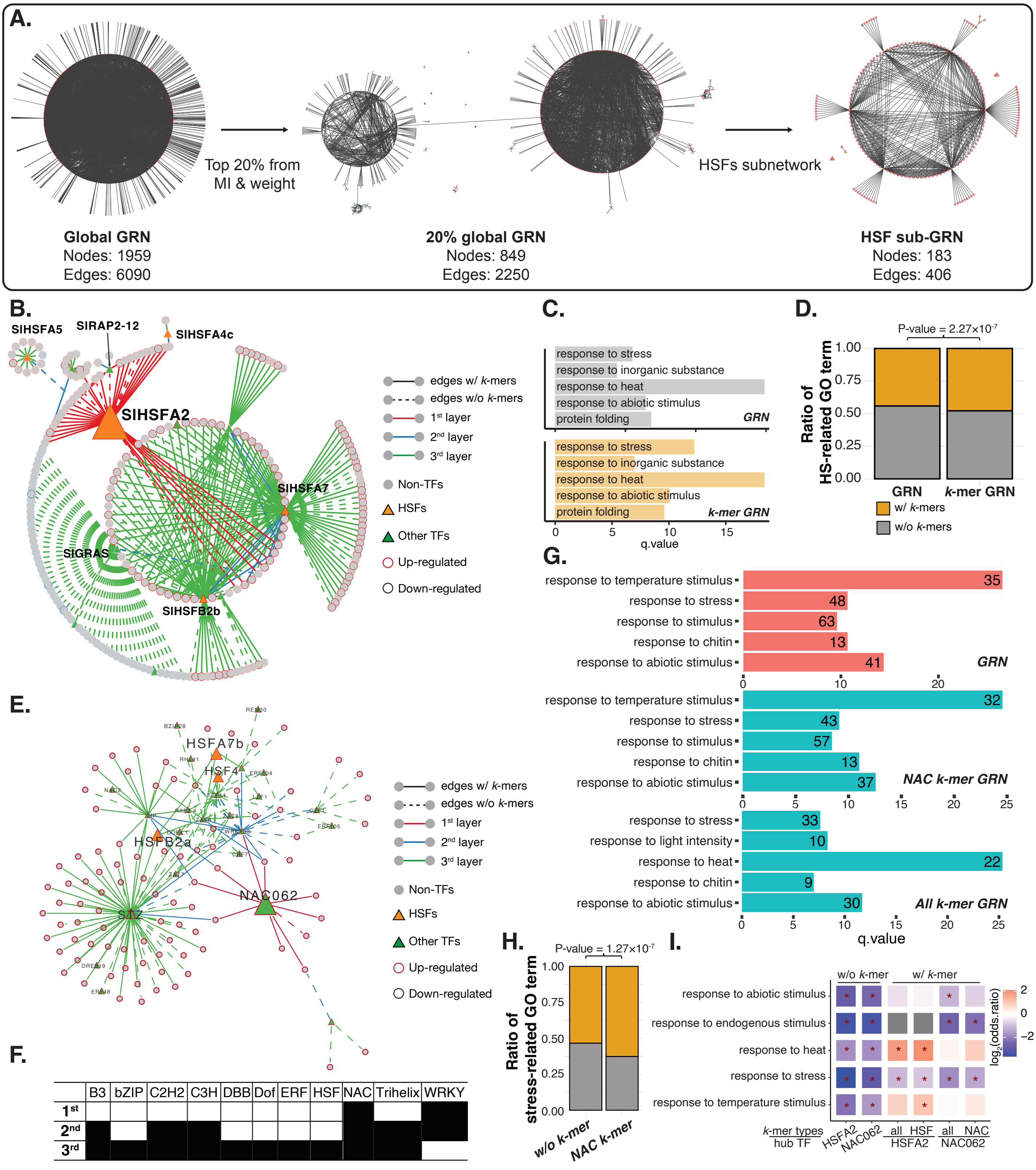

### Supplemental Figure 3

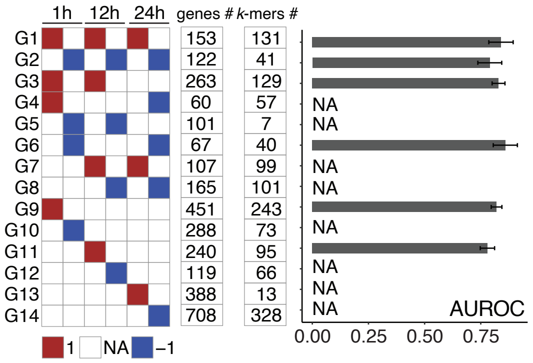
